# Can a short math video enhance the brain’s mathematical networks?

**DOI:** 10.1101/2022.08.09.503350

**Authors:** Marie Amalric, Pauline Roveyaz, Stanislas Dehaene

## Abstract

Many teaching websites, such as the Khan Academy, propose vivid videos illustrating a mathematical concept. Using fMRI, we asked whether watching such a video suffices to rapidly change the brain networks for mathematical knowledge. We capitalized on the finding that, when judging the truth of short spoken statements, distinct semantic regions activate depending on whether the statements bear on mathematical knowledge or on other domains of semantic knowledge. Here, participants answered such questions before and after watching a lively five-minute video which taught them the rudiments of a new domain. During the video, a distinct math-responsive network, comprising anterior intraparietal and inferior temporal nodes, showed inter-subject synchrony when viewing mathematics course rather than control courses in biology or law. However, this experience led to minimal subsequent changes in the activity of those domain-specific areas when answering questions on the same topics a few minutes later. All taught facts, whether mathematical or not, led to domain-general repetition enhancement, particularly prominent in the cuneus, posterior cingulate and posterior parietal cortices. We conclude that short videos do not suffice to induce a meaningful lasting change in the brain’s math-responsive network, but merely engage domain-general regions possibly involved in episodic short-term memory.

**Significance Statement:** Teaching mathematical concepts is difficult. To facilitate the comprehension and appeal of mathematics, several teaching websites provide vivid videos illustrating math concepts. Here, however, we show that merely watching such videos fails to improve the brain networks for mathematics. During the video itself, these networks are transiently engaged – but a few minutes later, when we ask questions about the taught concepts, performance is only minimally improved, and the participants engage generic regions thought to be involved in short-memory and language, rather than the targeted math-responsive regions. Brief video watching is therefore insufficient as a pedagogical device, probably because it misses ingredients such as teacher-pupil interactions, explicit teaching, active engagement, retrieval practice, repetition, and sleep.

## Introduction

Over the past decade, new modes of education have appeared. Massive on-line courses (MOOCs), educational video channels, computer-based gamified educational content, and other open online resources have simplified access to education for a large population. However, creating such digital education tools is not sufficient: their pedagogical impact must also be evaluated. The tools of cognitive neuroscience may help to formally evaluate them, in education settings as well as in lab settings (1). While previous studies have mostly used Randomized Control Trials to evaluate the amount of behavioral improvement compared to a classical pedagogical method (e.g., 2–6), here we show how functional magnetic resonance imaging (fMRI) can be used to track the brain mechanisms of such improvement.

In a pedagogical context, behavioral improvement is arguably the only thing that matters. However, neuroimaging can provide complementary insights about the underlying brain mechanisms and may sometimes strongly qualify the observed behavioral progress. For example, Yoncheva and collaborators (7) showed that, when adults learned to read a new alphabet using either a whole-word or a letter-based attentional focus, behavioral performance for the trained materials improved similarly in both groups, but the underlying change in brain activity was radically different: whole-word training lead to a right-hemispheric bias towards inappropriate circuits distant from the normal left-hemisphere regions for language and reading. In a similar vein, Delazer and collaborators (8) showed that learning a new arithmetic operation by drill versus by strategy elicits different brain activity in young adults. Functional MRI is now readily accessible to school-age children and college students, and has been used to track the fast brain changes underlying the acquisition of reading (9, 10) or arithmetic (11), as well as to evaluate the brain mechanisms underlying the superior performance of teaching strategies such as retrieval practice and creative mathematical reasoning (12).

While these studies mostly concentrate on skill acquisition, an even more relevant question from a pedagogical standpoint, indeed one of the most difficult and least understood ones, is how to teach the comprehension of a new domain. In the present study, we make one step in this direction by evaluating the neural correlates underlying the acquisition of curricular knowledge in mathematics and in non-mathematical domains. Neuroscience can now start investigating this issue thanks to advances in mapping the distributed brain networks underlying semantic knowledge (13–16). Different types of knowledge, for instance about actions, people, places, animals, or objects, appear to be broadly distributed in partially dissociable brain regions (16–19). In particular, mathematical knowledge, including but not restricted to the knowledge of numbers, can be dissociated from other forms of knowledge (20–22) and relies on a set of regions that respond more to mathematical statements than to other well-matched non-mathematical statements, for instance in geography or history (23–26). In particular, the intraparietal sulcus of both hemispheres, which has long been implicated in mental arithmetic (27, 28) and its acquisition (11, 29), is partially tuned to specific numbers (30–33) but also responds to more abstract mathematical concepts in professional mathematicians (24). So does the more recently discovered posterior inferior temporal gyrus, which was first thought to process the visual form of Arabic numerals (34), but in fact responds bilaterally during mental arithmetic and abstract math (24, 25, 35). Finally, large sectors of lateral prefrontal cortex, dorsal to language-related activations, are also activated by numbers and other math concepts (13, 14, 23–25).

By comparing expert mathematicians versus professors of other disciplines, our previous study suggested that this “math-responsive network” is capable of integrating additional mathematical knowledge (24). When listening to the very same statements, only expert mathematicians activated the math-responsive network when asked to determine the truth value of statements related to abstract math concepts that they had mastered for many years, and showed additional activity when the statements were meaningful rather than meaningless. Thus, the experts managed to integrate, within the same set of areas already engaged in arithmetic in all humans, additional mathematical knowledge.

Here we capitalize on these findings to ask whether we could visualize the rapid integration of new concepts to the math-responsive network. In the course of learning, a statement which is initially meaningless, because it involves unknown concepts, should become meaningful once those concepts are learned, and should join its appropriate place within the brain’s semantic networks, depending on the category of knowledge involved.

During an fMRI exam, we used four 5-minute pedagogical videos to teach new concepts to 21 freshmen math students from Parisian universities. Two of the videos introduced math topics that are typically introduced only much later, during the senior year (measure theory and stochastic processes). Two other videos served as controls and introduced non-math concepts (plant biology and property law). Learning was assessed by pre- and post-tests conducted inside the fMRI. In these tests, participants were asked to judge, as fast as they could, the veracity of spoken math and non-math statements (Figure 1). Half of the statements directly probed participants’ understanding of the newly taught concepts, while the other half bore upon either already known notions (taught during high school), or completely unknown notions (untaught). Thus, the design was a 2×2×3 factorial design with factors of domain (math versus control), time point (pre versus post video training), and content (known, taught, or untaught statements) (see Figure 1 and Methods for details).

**Fig 1.**
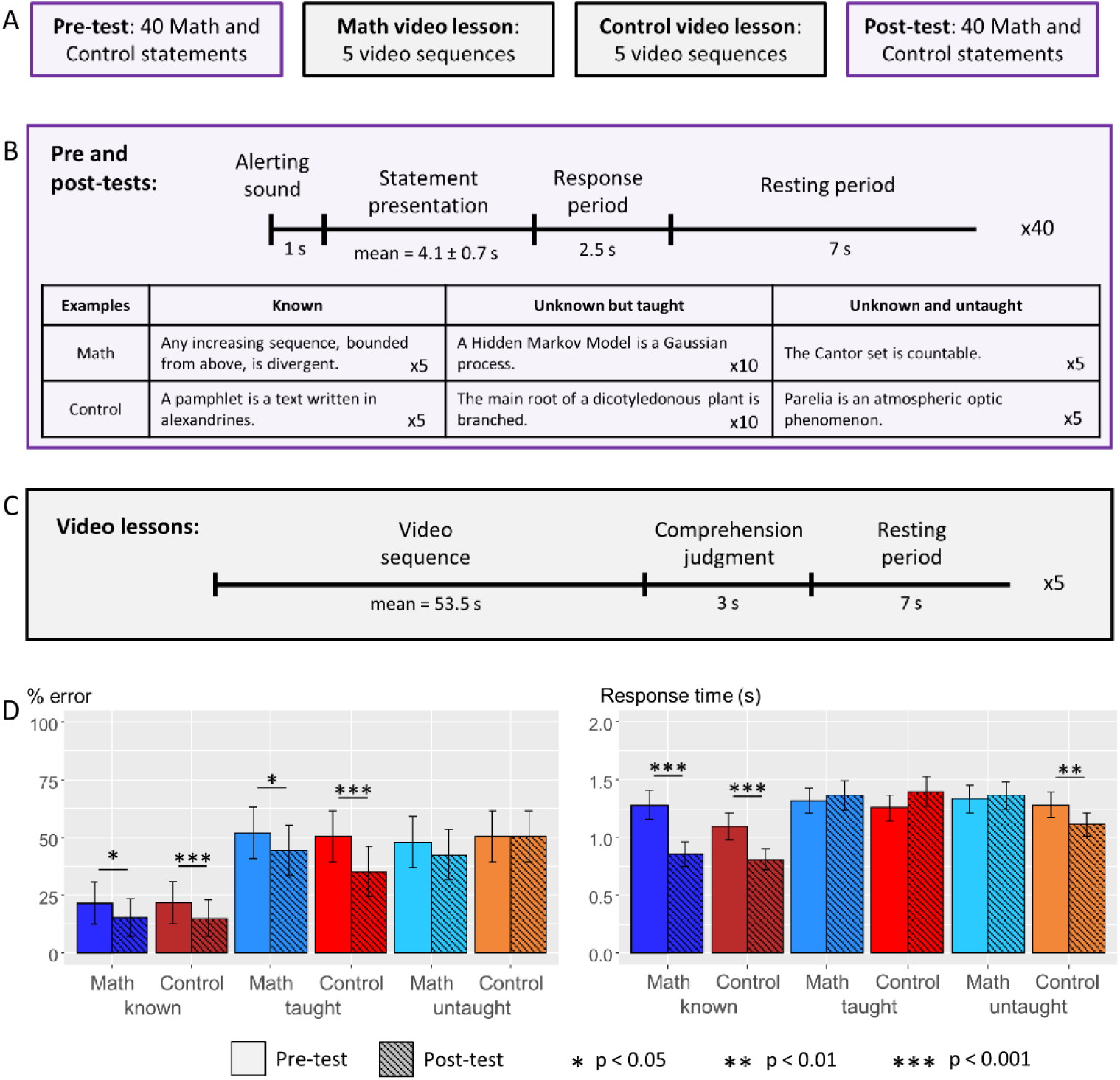
Protocol and behavioral results. (A) The fMRI experiment comprised 2 sessions, each composed of a pre-test, a math and a control video lesson in random order, and a post-test. (B) Structure of the pre- and post-test runs: 40 statements were presented orally, each followed by a short response period and rest period. The table shows examples of statements for each category: math/control x known/taught/untaught. (C) Structure of the video lessons: each lesson comprised 5 one-minute video sequences, each followed by a rapid introspective evaluation of the participant’s comprehension level. (D) Evolution of performance from pre-tests to post-tests, as assessed by error rates (left) and mean response times (right). Error bars indicate one standard error of the mean. Stars indicate the significance level of the difference between pre- and post-tests.

We had three main predictions. (1) For known facts, the contrast between math versus control statements should replicate earlier findings on the existence of a math-responsive network (23–26); (2) Repeating those known statements a second time should lead to repetition suppression, a reduction in fMRI signals in the relevant areas; but repeating the unknown statements after the corresponding concepts were taught in the video should, on the contrary, lead to repetition enhancement (36–38). The location of this increase in fMRI signal, which should occur for taught but not for untaught concepts, should indicate which brain circuits are responsible for acquiring this new knowledge. If short videos are pedagogically efficient, taught statements should yield repetition enhancement in their respective math-responsive versus general-knowledge circuits. (3) During the educational videos themselves, fMRI signals should exhibit inter-subject correlations (39–41), again targeting the relevant semantic networks. The magnitude of such inter-subject synchrony may be a marker of the amount of learning, as measured by both behavior and fMRI.

## Results

### Behavioral learning effect

We first verified that, in the pretests, participants responded way above chance to the supposedly known statements (math: 21 ± 9.1 % errors in 1.3 ± 0.13 secs; control: 22 ± 9.3 % errors in 1.1 ± 0.12 secs), but performed at chance on all the other statements (math taught: 52 ± 11 % errors in 1.3 ± 0.11 secs; math untaught: 48 ± 11 % errors in 1.3 ± 0.12 secs; control taught: 51 ± 11 % errors in 1.3 ± 0.11 secs; control untaught: 51 ± 11 % errors in 1.3 ± 0.11 secs) (Fig 1). Participants made fewer errors when judging the truth value of known statements (F(2,40) = 53.4, p < 10^−11^) and also answered faster (F(2,40) = 4.65, p = 0.015).

Participants’ performance improved after watching the video lessons on statements related to the taught concepts (math: −8.15% errors, 0.046 secs longer; control: −16.5% errors, 0.15 secs longer; error rate: both p*s* < 0.05, FDR corrected for multiple comparisons; response time: n.s.; Fig 1). Performance also improved slightly with known statements (math: −6.24% errors, 0.44 secs faster; control: −6.88% errors, 0.28 secs faster; all p*s* < 0.05; Fig 1), but not with unknown statements (math: −4.94% errors, n.s, 0.031 secs longer, n.s.; control: +0.91% errors, n.s., 0.17 secs faster, p < 0.01; Fig 1).

During the videos, once per minute, participants rated their comprehension on a 1-4 scale. When transformed to a 0-100 scale, average comprehension reached 69.7% in math and 72.3% in control videos, a non-significant difference (t(20) = 0.93). There was a positive correlation, across participants, between comprehension ratings and performance improvement (math: r = 0.26; control: r = 0.50), though this effect reached significance only for the control statements (p = 0.02).

### Math semantics is systematically processed in a distinct set of brain regions

We first tested prediction 1: different network should be activated by math versus control statements related to known concepts. For this analysis, known statements from both pre- and post-tests were pooled. This contrast revealed the classical set of math-responsive brain regions, including bilateral anterior and posterior intraparietal sulci (aIPS and pIPS), bilateral posterior inferior temporal gyri (pITG), bilateral superior frontal gyrus (SFG), and the left middle prefrontal gyrus (Fig 2A, Table S1), at locations similar to the ones found in our previous research (24, 25). The converse contrast of control versus math known statements showed activity in the left temporal pole and inferior frontal gyrus, bilateral fusiform gyrus, postcentral gyrus, and middle occipital gyrus.

**Fig 2.**
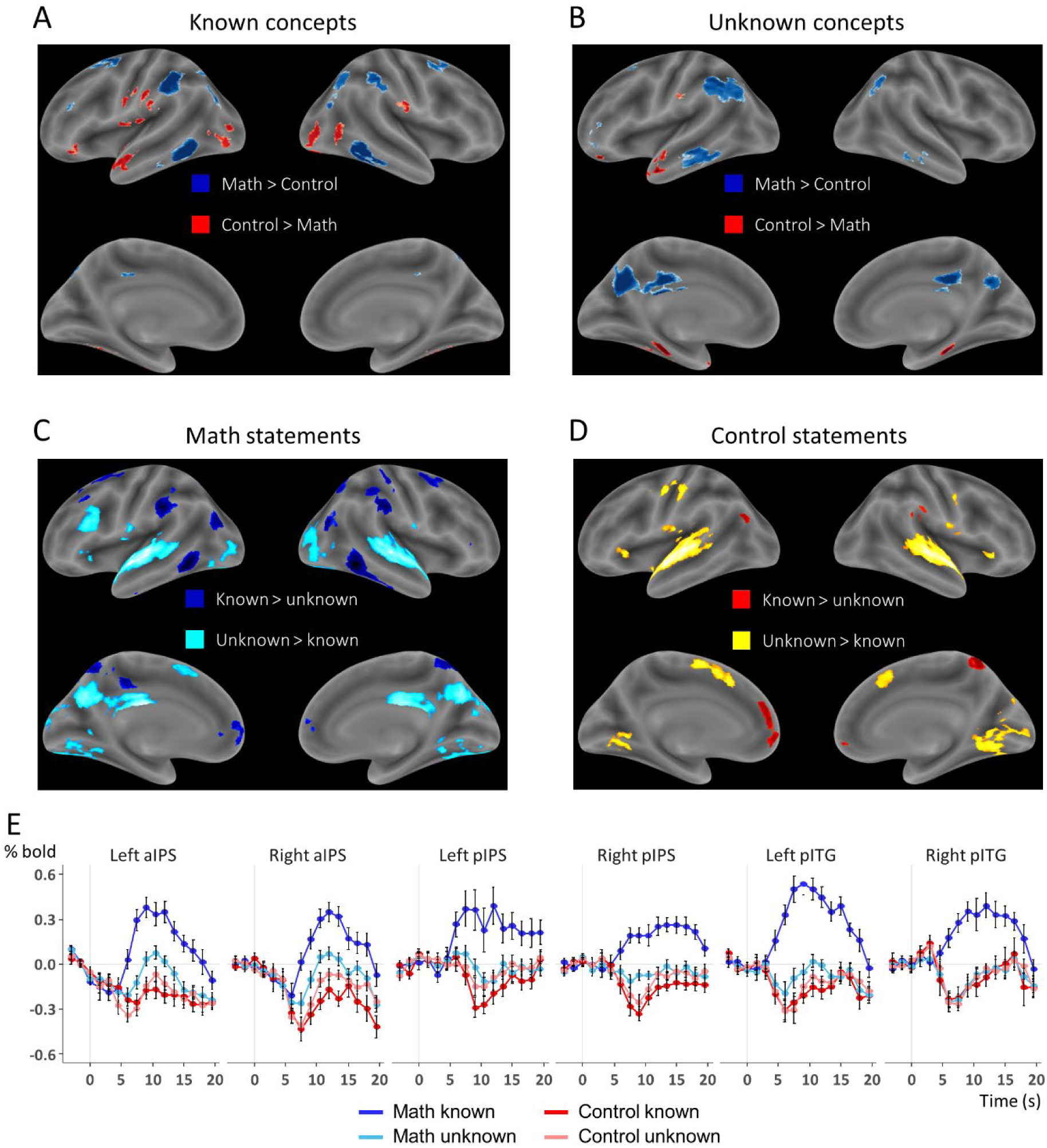
Distinct brain networks for mathematical versus general knowledge. Whole-brain inflated maps showing the comparison of activity elicited by (A) known math statements more than known control statements in blue, and the opposite contrast in red; (B) unknown math statements more than unknown control statements in blue, and the opposite contrast in red; (C) known math statements more than unknown math statements in dark blue, and the opposite contrast in cyan; (D) known control statements more than unknown control statements in red, and the opposite contrast in yellow. All maps are thresholded at p < 0.001 uncorrected voxel-wise, and p < 0.05 FDR-corrected cluster-wise. (E) Time course of the BOLD signal for each category of statement in representative brain areas of the network responsive to mathematics: left and right anterior and posterior IPS (aIPS and pIPS), and left and right posterior inferior temporal gyrus (pITG).

We next asked whether the unknown concepts, although unmastered by the participants, were already mapped onto the same differentiated circuits for math vs control concepts, as they partially made use of the same math vocabulary. We thus compared math versus control statements involving unknown concepts (pooling together both taught and untaught concepts in pre-tests, as well as untaught concepts in post-tests). Although the differences were weak, there was a slightly greater activation for math statements compared to control statements mostly in the left hemisphere in the angular gyrus (AG) extending through the inferior parietal lobule (IPL), and in the inferior temporal gyrus (ITG), as well as in the posterior cingulate cortex (PCC) and cuneus along the brain midline (Fig 2B, Table S1). While the parietal and inferior temporal sites were close to the math-responsive network, they only partially overlapped.

The direct contrast of known versus unknown math concepts (Fig 2C) yielded a map similar to the one comparing known math versus control statements, again revealing the classical math-responsive network composed of bilateral aIPS, bilateral pITG, and various middle and superior frontal sites (Fig 2C, Table S2). In the converse direction, unknown math concepts induced more activity than known math concepts in the bilateral superior temporal sulcus (STS), the bilateral lingual and middle occipital gyri, the left inferior frontal gyrus, as well as in the PCC and cuneus. These regions are outside of the math-responsive network, and partially overlap with classical areas for spoken or written sentence processing, thus suggesting that sentences comprising unknown words may have induced a greater amount of difficulty within language comprehension networks. Indeed, these results, particularly the greater activation of the bilateral superior temporal regions and lingual areas, were partially replicated when contrasting unknown versus known control concepts (Fig 2D, Table S2).

These results are illustrated in Fig 2E by plots of the average timecourse of the neural response within the six main math-responsive regions revealed by the contrast of math versus control statements (bilateral aIPS, pIPS, and pITG). In those regions, following the auditory presentation of the statement, the BOLD signal systematically rose to known math concepts, and to a much lesser extent to unknown math concepts in both aIPS, but no activation or even a deactivation was seen in all other cases. Taken together, these results suggest that known math statements differed from all other statements in their capacity to strongly activate the math-responsive network (23–26).

### Learning effect due to exposure to the videos

We then evaluated the main goal of our experiment, i.e., whether the learning sessions affected brain activity by channeling the taught math statements towards the math-responsive network. To do so, we compared the neural activity elicited by post-test versus pre-test presentations of the same math statements, before and after the relevant concepts were taught. However, this contrast did not reveal the expected math-responsive network. Rather, repetition enhancement was seen in the posterior mesial part of the brain, i.e., in the cuneus/precuneus and posterior cingulate cortex accompanied by a large swath of activity in the left posterior inferior parietal cortex/angular gyrus, the anterior part of the middle frontal gyrus, and bilateral caudate nuclei (Table S3).

Importantly, this result was not specific to math. On the contrary, a very similar network including the cuneus/precuneus, posterior cingulate cortex, left inferior parietal lobule/angular gyrus, middle frontal sites, and bilateral caudate nuclei was observed for the pre- versus post-contrast with taught control statements (Table S3). Indeed, no region showed a significant interaction (post – pre) X (math – control) for taught concepts.

Because activation could change across time or sentence repetition without implying a learning effect, we replicated the above analysis using two more specific contrasts: C1 = (post – pre) X (taught – untaught), and C2 = (post – pre) X [taught – (known & untaught)]. The results were similar, consistently revealing activity in the caudate nuclei, left inferior parietal lobule/angular gyrus, cuneus/precuneus and middle frontal gyrus (Table S3). Again, there was no difference between math and control, as no region showed significant interactions C1 X (math – control) or C2 X (math – control). Figure 3A shows the main effect of repetition enhancement imputable to learning, pooling over both math and control concepts (contrast C2).

**Fig 3.**
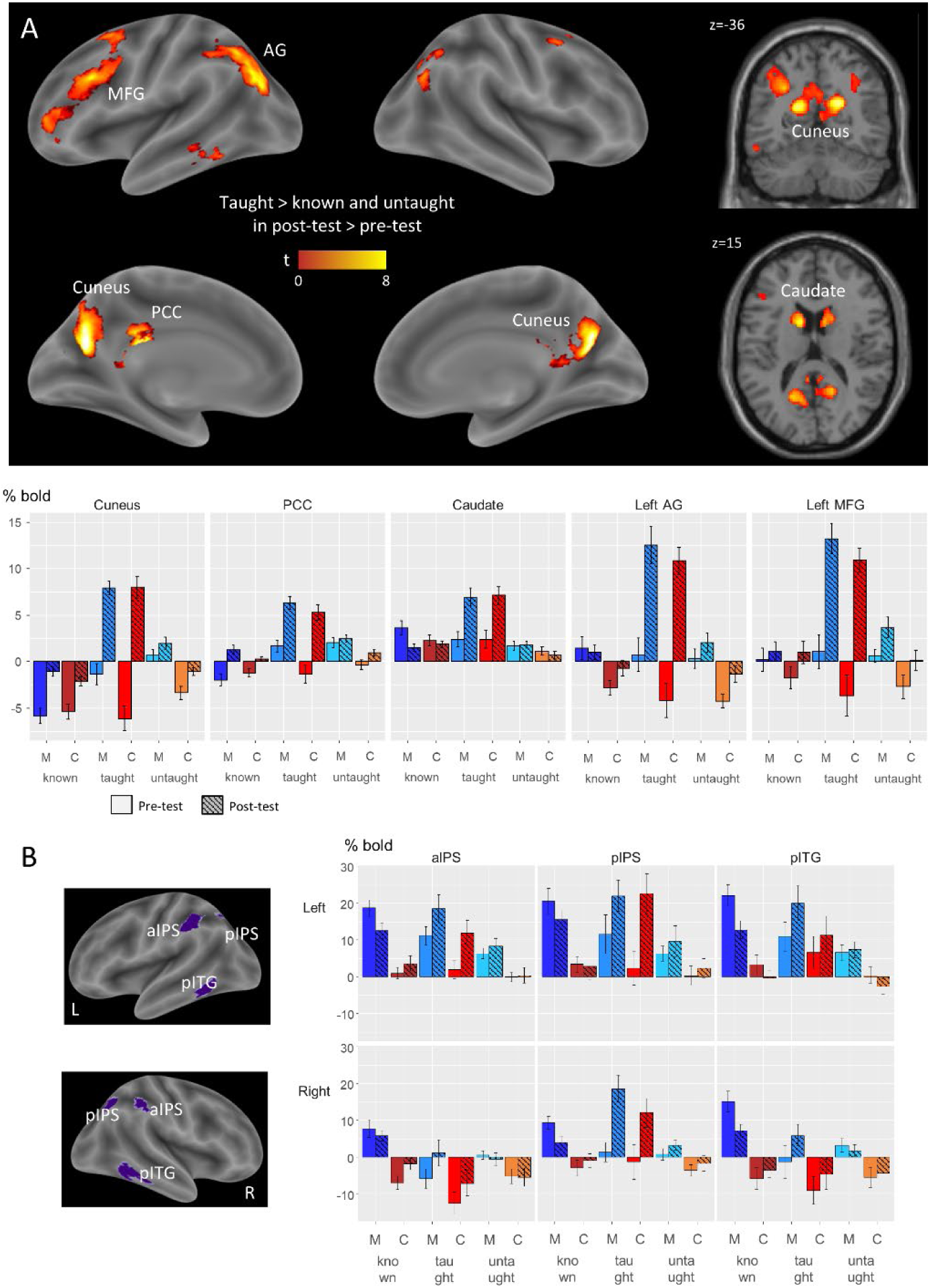
Effects of learning at the brain level. (A) Interaction effect indicating significantly greater activity during post-tests than during pre-tests, for taught statements more than for known and untaught statements, pooling across math and control statements (voxel-wise p < 0.001 uncorrected, cluster-wise p < 0.05 with FDR correction). Histograms show the mean beta estimates for each category of statements in both pre- and post-tests, extracted from the principal clusters of activation. (B) Mean beta estimates for each category of statements in both pre- and post-tests (right), extracted from the six main math-related regions of interest, displayed on an inflated brain (left).

The histograms of figure 3A illustrate the details of these effects. A small amount of repetition enhancement occurred even to known statements in some regions (posterior cingulate cortex, cuneus/precuneus, and middle frontal gyrus). However, all regions showed a much larger increase in activation selectively for taught statements after viewing the video. Furthermore, such learning-induced repetition enhancement was identical for math and non-math, and therefore unselective for contents.

The regions exhibiting an increase of activation after viewing the videos partially overlap with regions involved in episodic memory and language processing, but not within the math-responsive brain network. We further confirmed that the newly taught concepts did not trigger the activation of the math-responsive network thanks to a sensitive analysis using subject-specific voxels within the 6 main math-related regions-of-interest (ROIs; Fig 3B). Voxels were identified using the contrast for simple calculation versus sentence processing from a short localizer completed at the end of the main fMRI experiment. We then conducted an ANOVA with hemisphere, region (aIPS, pIPS, or pITG), content type (math or control nonmath), statement category (know, taught, or untaught), and test phase (pre- or post-) as within-subject factors. We observed generally greater activation in the left hemisphere compared to the right hemisphere (F(1,19) = 35.1, p < 2.10^−5^), for math statements compared to control nonmath statements (F(1,19) = 29.4, p < 4.10^−5^). We also found a main effect of the statement category (F(2,38) = 7.61, p = 0.002), but no effect of the region of interest (F(2,38) = 1.97, p = 0.153).

Turning to the analysis of the effect of learning, we found repetition suppression for known math statements (F(1,19) = 17.4, p < 0.001), no significant difference between pre- and post-tests for untaught concepts (F(1,19) = 0.357, p = 0.557), and a clear repetition enhancement for both taught math and taught control concepts alike (Math: F(1,19) = 11.6; Control: F(1,19) = 11.7; both ps < 0.003). Indeed, the interaction (Math – Control) X (Taught – Untaught) X (Pre - Post) was not significant (F(1,19) = 0.042, p = 0.839). Thus, freshly learned math concepts yielded enhanced activity in the math-responsive brain regions, but in the same proportion as freshly learned control concepts.

### Inter-subject correlations during exposure to the videos

The above results indicated that a short period of learning was not sufficient to selectively channel the taught math concepts towards the math-responsive network, and instead mobilized the precuneus and related regions not specifically involved in mathematics (Fig 3). To clarify what happened during the learning process, we investigated participants’ neural responses during the video lessons themselves. Because videos are complex stimuli that involve rich visual and auditory inputs combined in elaborate narratives, we analyzed neural activity during video watching by calculating inter-subject correlations (ISC), a data-driven analysis technique that locates any brain region reliably entrained by the videos in a systematic way across participants.

Figure 4A shows the comparison of ISC during math versus control videos. Relative to control videos, math videos elicited more synchronous activity among participants in bilateral IPS and right pITG, as well as at various frontal sites of the right hemisphere (Table S4). The converse contrast revealed synchronous activity along the superior temporal sulcus of both hemispheres, in the primary visual cortex, in the bilateral fusiform gyrus, and the left inferior frontal gyrus. Therefore, our videos were partially successful in engaging the appropriate semantic networks, in a distinct manner for math versus control videos, in agreement with previous results on the capacity of brief pedagogical videos to activate those brain systems (39, 42, 43).

**Fig 4.**
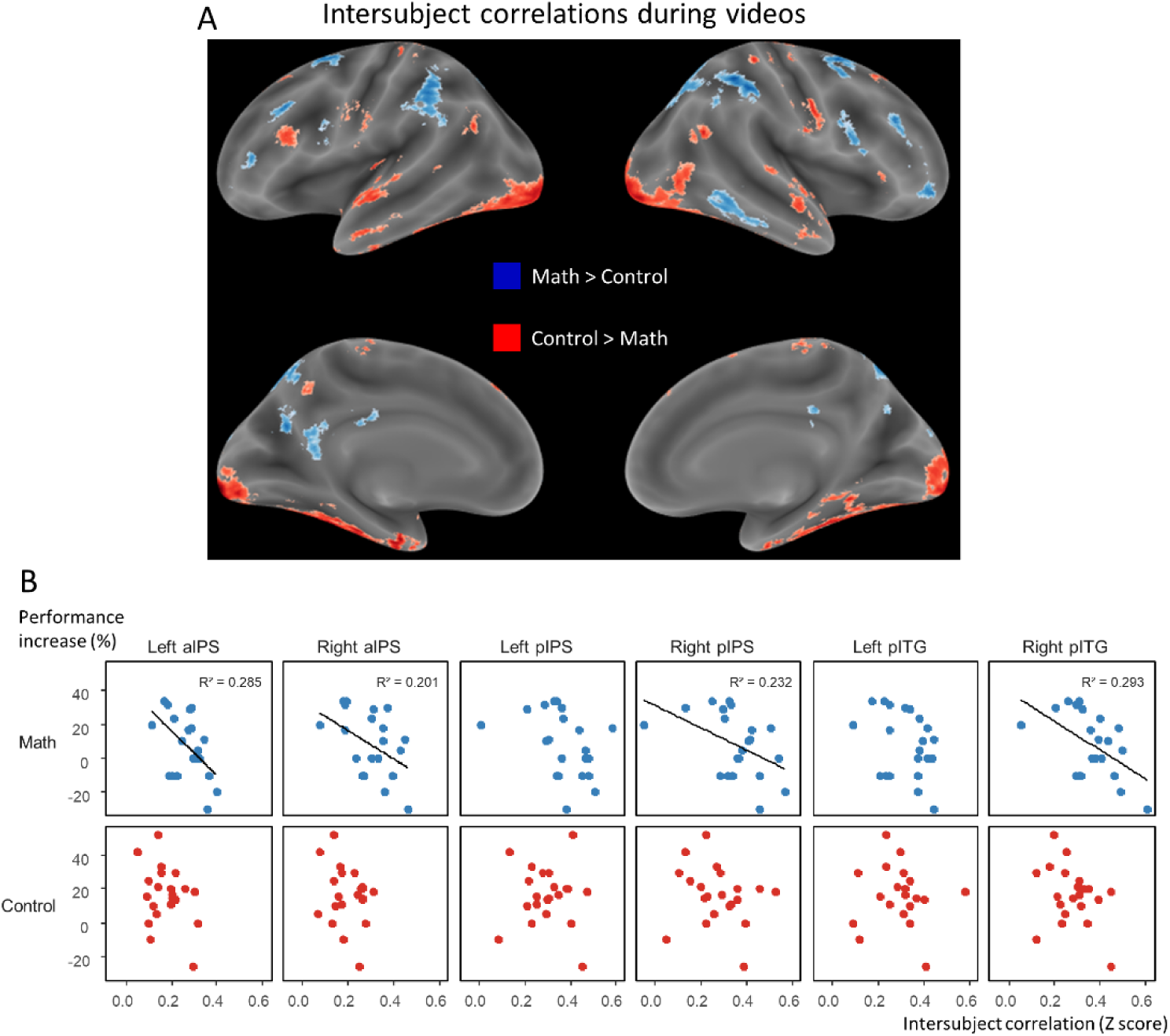
Analysis of brain activity during the video lessons. (A) Whole-brain inflated maps showing the regions with temporally correlated activation across participants (ISC) during math video lessons more than during control video lessons in blue, and the opposite contrast in red (voxelwise p < 0.001 uncorrected, clusterwise p < 0.05 FDR-corrected). (B) Correlation between the mean ISC value extracted from the six main math-related regions of interest, and the performance increase from pre-to post-tests across participants, separately for math (top row) and control (bottom row). We observed negative correlations between ISC during math video lessons and the performance increase on taught math statements in most regions of interest. In the same regions, there was no significant correlation between ISC during the control nonmath video lessons and the performance increase on taught control statements.

Previous research suggested that children’s neural maturity during video watching, as assessed by the similarity of their activation patterns to those of adults, predicted their performance in arithmetic and letter knowledge (39). Here, we similarly asked whether the similarity of a person’s math-related brain responses to the group, during video watching, reflected the extent to which they learned from it. We thus investigated whether inter-individual differences in learning (i.e., performance increase from pre-to post-tests) were related to inter-individual differences in the strength of neural response correlation, in the math-responsive regions, between each individual and the rest of the group while watching the video lessons.

Contrary to what we expected, we found significant *inverse* correlations in almost all pre-defined math-related regions of interest (Left aIPS: r = −0.534; Right aIPS: r = −0.448; Left pIPS: r = −0.400; Right pIPS: r = −0.481; Left pITG: r = −0.341; Right pITG: r = −0.541; reaching significance in all regions but both left pIPS and pITG) (Fig 4B). The higher an individual ISC during the math video lessons, the smaller his or her increase in performance on taught math concepts. No such pattern was found between individual ISC evaluated during the control video lessons and the increase in performance on taught control concepts. Inspired by previous findings (39, 44), we also explored which regions of the brain exhibited a positive correlation between intersubject synchrony and the amount of learning. To do so, the Pearson correlation coefficient between ISC and performance increase was computed across participants in each voxel of the brain, separately for the math and control videos. We found positive correlations in small clusters outside the math-responsive network, particularly in the para-hippocampal gyrus (bilateral for math videos, [−30, −18, −24], r=0.725, and [18, −16, −24], r=0.672; right only for control videos, [26, 7, −28], r=0.656), but none survived a voxel-wise threshold of p < 0.001. These results suggest that, in the present context of short-term video learning, the synchronization of neural responses with the rest of the group, within the math-responsive network, primarily entertains a negative relation to learning.

## Discussion

Our goal was to the study the impact of a short video lesson on the brain’s semantic networks. To this aim, we scanned participants with fMRI while they answered simple true/false questions on mathematical and non-mathematical topics, before and after being exposed to a 5-minute educational video. Learning did occur, as evidenced by changes in both behavioral accuracy and brain activity evoked by statements in the taught domain, but not in previously known or in untaught domains. However, activation did not increase in areas encoding domain-specific semantic knowledge, but only in a domain-general network prominently including the cuneus/precuneus, angular gyrus and bilateral caudate, and indifferently so for math and non-math statements. We now discuss what these results imply for brain organization, pedagogy, and video-based learning.

Our work was predicated upon the dissociation, reported in several prior studies, between brain regions involved in the representation of mathematical facts and in other domains of knowledge (22– 28). It is therefore important that we replicated these observations in two different ways. First, for facts that participants already knew before the experiment, we observed a clear separation between those two sets of regions (figure 2A). As in previous work, mathematical statements, compared to control non-mathematical statements induced greater activity in bilateral anterior intraparietal and posterior inferotemporal regions, while control non-mathematical statements induced greater activity in other areas classically involved in high-level semantic integration and memory, notably the bilateral temporal poles (15, 45, 46). The present experiment also included a novel control, the contrast of known versus unknown facts. Although less diagnostic, perhaps because participants searched their semantic memory when faced with unknown facts, this contrast evidenced a bilateral anterior intraparietal as well as right ventral temporal activation specific to known mathematical facts (figure 2C). Those within-subject results, obtained in undergraduate math students, concur with a prior between-subjects comparison of mathematicians versus non-mathematicians (24). Together, they indicate that those regions, while systematically involved in encoding numbers and simple arithmetic (30–33), are transformed by mathematical education and end up encoding more complex, non-numerical learned mathematical facts.

Second, similar observations were made using a different method, inter-subject correlations (ISC) in brain activity while participants watched the same pedagogical movies (figure 4). During math movies, compared to control non-math movies, inter-subject synchronization was stronger in bilateral anterior intraparietal and right ventral temporal areas, i.e. the same regions that were previously found to be math-responsive and selective for known math facts (compare figure 4 and figure 2). In the converse direction, non-math movies tended to evoke greater synchrony in a more distributed set of areas, again including left and right anterior temporal regions. In past studies, fMRI inter-subject synchrony in naturalistic settings (40, 41, 44) was found useful to evaluate, not only the activation of sensory areas driven by specific stimuli such as faces, but also of more abstract, semantic cortical regions, for instance those shared between movie watching and story listening (47), spoken and written version of the same story (48), bilingual presentations of the same story (49), or between listening to a story and producing it (50). When people diverged in their comprehension of an ambiguous story, their ISCs reflected which meaning they grasped (47, 51). Thus, ISC seems ideal to track whether and how students understand a pedagogical lesson. Indeed, several recent studies have shown that ISC can track teacher-student dynamics using fMRI (44, 52), but also the lighter techniques of functional near-infrared spectroscopy (53) and electro-encephalography (54, 55). Furthermore, a previous fMRI study showed content-selective ISC in preschoolers watching Sesame Street videos, with intraparietal cortex showing larger ISC during number-related segments of the program (39). The present study confirms this finding: ISC varied with the content of the videos and concentrated in the bilateral intraparietal sulcus and other math-responsive areas during math lessons (Figure 4).

Following the video, as predicted, we found repetition enhancement, i.e. an increase in fMRI signals evoked specifically by statements bearing on the learned materials. Thus, our results confirm that repetition enhancement can serve as a marker of learning (36–38). Nevertheless, the impact of such video training on subsequent judgements differed from our predictions in two ways. First, learned statements failed to evoke content-specific activation: no difference was found by taught math versus taught non-math statements. Second, the regions where learning occurred (as evidenced by larger repetition enhancement for taught concepts than for known or untaught ones) – cuneus, posterior cingulate, caudate, angular gyrus and middle frontal gyrus – failed to be strongly activated by previously known statements (Figure 3A). Thus, immediately following learning, the participants had not yet “routed” the learned facts to the relevant content-specific brain areas, and instead, held them in a distinct network. Attributing a function to such a diverse set of regions is difficult, given the well-known pitfalls of “reverse inference”, i.e. the impossibility of determining of cognitive processes from brain imaging data (56, 57). However, we note that part of this network (angular gyrus, precuneus, PCC) is active during high-level language processing and participates in a slow semantic integration system which is thought to store the meanings of passages or movies in episodic memory (58, 59) and replay them in imagination (60). Our results concur with a previous study where learning to solve mathematical problems also led to short-term repetition enhancement in posterior cingulate, precuneus and retrosplenial cortex as participants repeatedly practiced solving the same problems (37). Together, these observations suggest a putative interpretation of our results: learning may be initially biased to domain-general short-term, episodic memory systems. The videos seem to have failed to induce a deep understanding of mathematical meaning. Instead, the participants would have answered the questions, at the post-video stage, by evoking a rote literal memory of the movies (verbal or visual). Indeed, such a shallow strategy is perhaps the only reasonable or accessible one when a student is bombarded with 5 minutes of new concepts and vocabulary.

This interpretation may also provide an explanation for the surprising finding of a negative correlation, across participants, between ISC in the math-responsive network, and the amount of learning. In their study of Sesame Street videos, Cantlon and Li (39) demonstrated that the amount of synchrony with adults in intraparietal cortex was a positive predictor of math performance. Likewise, Hasson et al. (44) observed that enhanced ISC was a positive predictor of episodic memory 3 weeks later. The present conditions are quite different, however, and the negative correlation may be due to the fact that it is essentially impossible to integrate so many new facts into the mathematical semantic network in just five minutes. Therefore, the participants who succeed in answering some of our questions after watching the videos are not those who tried to deeply understand the math, but those who merely stored some of the taught materials using a domain-general episodic memory network. In agreement with this interpretation, we found a trend toward positive correlations across participants between ISC and the amount of learning in the parahippocampal gyrus, i.e. regions similar to those identified by Hasson et al. (44) and linked to episodic memory retrieval (61–64).

While this specific interpretation is clearly speculative and will require further testing, one conclusion is clear: an entertaining five-minute pedagogical video fails to teach mathematical facts at a deep level. Furthermore, as we discussed in the introduction, behavioral evidence of learning may be insufficient, and brain imaging may provide a complementary viewpoint by showing whether or not the relevant brain networks are activated (7). Our results suggest that much caution is needed before spreading broadly the short videos that the Khan Academy and many other websites provide. Undoubtedly, those videos may be useful to convey intuitions, visualizations, and enthusiasm for math, but more experimentation is needed to probe the actual, operational knowledge that they leave in the viewer’s memory. The literature on the scientific foundations of learning (65–68) suggest that brief video watching lacks several ingredients that could be crucial for efficient learning, including eye-to-eye pedagogical teacher-pupil interactions (69), explicit and direct instruction (70), active pupil engagement (12, 71), retrieval practice (72), spaced repetition (73), and sleep-dependent consolidation (74). In the future, the present fMRI design could be used to test those factors.

## Materials and Methods

### Participants

21 math major students, all in their freshman or sophomore year in math, took part in this study (14 male, 7 female, age range 18-23, mean = 20). They were recruited in various Parisian universities and preparatory classes. They were included in this experiment based on their answers to a questionnaire insuring that did not already know what was about to be taught but mastered the background knowledge mandatory to understand the lessons.

All participants gave written informed consent and were paid for their participation. The experiment was approved by the regional ethical committee for biomedical research (Comité de Protection des Personnes, Hôpital de Bicêtre).

### Procedure

The fMRI exam was divided into two consecutive sessions. Each session started with a knowledge pretest in which participants were asked to judge as fast as they could whether forty spoken math and control nonmath statements were true or false. Participants were then presented with one math and one nonmath pedagogical video lessons. The order of video lessons was counterbalanced between the two sessions (session1: measure theory followed by plant biology; session 2: property law followed by stochastic processes), and the session order was randomly assigned to participants. Each session ended with a knowledge post-test, consisting of the same forty spoken math and control nonmath statements that were presented during pre-test (Figure 1). After the fMRI exam, participants were asked, without time limit, to give their definitions of the concepts taught by the 4 video lessons they watched.

#### Test runs

Participants were presented with a total of 40 math statements and 40 control nonmath statements. In each session, 5 math and 5 control statements were related to familiar notions extracted from the high school math and French curricula; 10 math and 10 control statements were related to the concepts introduced in the video lessons; 5 math and 5 control statements dealt with completely unknown concepts. The 10 math concepts introduced during the fMRI exam were part of the typical senior year math curriculum: measure theory (Sets, Countability, σ-Algebra, Lebesgue measure, and Lebesgue integral) and stochastic processes (Brownian motion, stochastic, Gaussian, and Markov processes, and Hidden Markov Model). The 10 control nonmath concepts introduced in this experiment came from completely unrelated fields that also typically use very specific vocabulary: property law (legal person, estate, inheritance, usufruct, annuity) and plant biology (gymnosperm, angiosperm, dicotyledon, spermatophyte, graminoid).

Within each category of statements (known, taught, untaught), half were true, and half were false. Reference to numbers or to other mathematical concepts was purposely avoided. A complete list of statements, translated from the original French, is presented in SI.

All statements were recorded by a female native French speaker who was familiar with mathematical concepts. Statements from the different categories were matched in syntactic construction, length (mean number of syllables: math = 16.9; control = 17.2; t = − 0.58, p = 0.56, no significant difference between categories: F(5,74) = 0.61, p = 0.69), and duration (math = 3.28 s; control = 3.23 s; t = 0.80, p = 0.42, no significant difference between categories: F(5,74) = 1.25, p = 0.29).

Each session comprised two test runs, one before and one after the presentation of the videos, that both included the same 40 statements presented in random order. On screen, the only display was a fixation cross on a black background. The onset of each statement was announced by a beep and the fixation cross briefly turning to red. After the auditory presentation of a statement, the fixation cross turned to green, signaling that a response was expected. Participants were given 2.5 seconds to evaluate whether the statement was true or false, by pressing one of two corresponding buttons (held in the right hand). Each trial ended with a 7-seconds resting period (Figure 1).

#### Video runs

Each video run consisted of two video lessons separated by a 30-seconds resting period. Each video lesson was composed of five 53-seconds video sequences, each teaching one given concept, progressing from the most basic concept to the most complex one (Figure 1).

The video sequences were created using the application “Explain Everything”, by one of the authors, PR, who was a mathematics teacher. On a white background, they presented a combination of pictures, animations, keywords, and auditory explanations. Math and control video sequences were matched in duration (math = 55 s, control = 52 s, t=1.03, p = 0.32).

After each video sequence, participants were given 3 seconds to report their level of understanding, on a 1-4 scale, by pressing one of the four corresponding buttons they held in the right hand. Each sequence ended with a 7-seconds resting period. The only screen display was a black fixation cross on a white background.

### Localizer run

This 5-minutes run is described in detail elsewhere (75), and includes mental processing of simple subtraction problems presented visually or auditorily, contrasted to the processing of written or spoken sentences of equivalent duration and complexity. This contrast was used to select voxels activated by math processing at the individual level.

### fMRI data acquisition and analysis

We used a 3-Tesla whole body system (Siemens Trio) with a 64-channel head-coil and multiband imaging sequences [multiband factor = 3, Grappa factor = 0, 60 interleaved axial slices, 1.75-mm thickness and 1.75-mm isotropic in-plane resolution, repetition time (RT) = 1,810 ms, echo time (ET) = 30.4 ms].

Using SPM12 software, functional images were first realigned, normalized to the standard Montreal Neurological Institute (MNI) brain space, resampled to 1.5 mm voxel size, and spatially smoothed with an isotropic Gaussian filter of 2 mm FWHM.

For test runs, a two-level analysis was then implemented in SPM12 software. For each participant, fMRI images were high-pass filtered at 128 s, and time series were modeled by one regressor for each statement (kernel = sentence duration + mean response time). Regressors of non-interest included the six framewise displacement parameters extracted from previous realignment. We defined subject-specific contrasts for each category of statements - Math/Control x Known/Taught/Untaught x Pre-/Post-test – versus rest. These contrasts were then smoothed with an isotropic Gaussian filter of 5 mm FWHM and finally entered in a second-level whole-brain ANOVA with statement conditions as within-subject factors.

For video runs, we used the technique of inter-subject correlation (ISC), which consists in identifying brain regions where the response to a stimulus is correlated among participants (76).

The 6 framewise displacement parameters were regressed out, and a 100-second high-pass filter was applied. Rest periods at the beginning of each run were then trimmed, resulting in four video runs of 169, 158, 148, and 163 TRs each. Each run was finally z-scored (standardized to zero mean and unit variance) for each voxel. An average gray-matter mask was applied prior to performing any further analyses. The inter-subject correlation was then calculated using the leave-one-out approach: at each voxel, each subject’s time series (i.e., each run) was correlated with the average time series of all the other participants (76). This approach gave us four R-maps for each participant that were converted into z-maps using the Fischer transformation. At the group level, the z-maps of each task were entered in a repeated measure ANOVA. All brain activation results are reported with an uncorrected voxelwise threshold of p < 0.001, and an FDR-corrected clusterwise threshold of p < 0.05.

## Acknowledgments

We thank the NeuroSpin teams for technical support, and Paul-Henri Icher for help in data collection. This research was supported by INSERM, CEA, Collège de France, the Bettencourt-Schueller Foundation, and a Marie Curie fellowship to M.A.

---

**Table S1.** MNI coordinates and t values of the main activation peaks for the contrasts of known math > control statements, known control > math statements, unknown math > control statements, and unknown control > math statements, extracted from maps thresholded at voxel-wise p < 0.001 uncorrected, and cluster-wise p < 0.05 FDR-corrected.

| Cortical Region | Hemisphere | MNI Peak Coordinates |  |  | t-value |
| --- | --- | --- | --- | --- | --- |
|  |  | x | y | z |  |
| Known Math > Control |  |  |  |  |  |
| Inferior Temporal | Left | -56 | -52 | -15 | 9.813 |
| Inferior Temporal | Right | 57 | -55 | -12 | 8.104 |
| Inferior Parietal | Left | -38 | -42 | 38 | 6.901 |
| Inferior Parietal | Right | 50 | -39 | 50 | 6.209 |
| Superior Parietal | Left | -18 | -78 | 49 | 5.349 |
| Superior Frontal | Left | -23 | 7 | 70 | 6.384 |
| Superior Frontal | Right | 29 | 13 | 56 | 6.737 |
| Middle Frontal | Left | -45 | 38 | 25 | 4.526 |
| Middle Occipital | Left | -38 | -82 | 32 | 4.721 |
| Middle Occipital | Right | 42 | -76 | 28 | 4.747 |
| Angular Gyrus | Right | 33 | -67 | 44 | 4.379 |
| Precuneus | Right | 11 | -69 | 59 | 4.076 |
| Middle Cingulate | Left | -3 | -30 | 43 | 4.543 |
| Known Control > Math |  |  |  |  |  |
| Inferior Frontal p. Orb | Left | -42 | 31 | -12 | 4.495 |
| Temporal Pole | Left | -42 | 14 | -25 | 4.751 |
| Middle Temporal | Right | 42 | -63 | 4 | 4.204 |
| Postcentral Gyrus | Left | -54 | -15 | 31 | 4.578 |
| Rolandic Opercularis | Left | -51 | -1 | 14 | 4.583 |
| Fusiform Gyrus | Left | -35 | -36 | -21 | 3.194 |
| Fusiform Gyrus | Right | 41 | -43 | -19 | 4.278 |
| Middle Occipital | Left | -38 | -66 | 5 | 4.731 |
| Cerebelum 6 | Left | -27 | -57 | -22 | 5.096 |
| Cerebelum 6 | Right | 32 | -61 | -22 | 3.612 |
| Unknown Math > Control |  |  |  |  |  |
| Inferior Parietal | Left | -42 | -45 | 37 | 5.524 |
| Angular Gyrus | Left | -44 | -63 | 50 | 4.711 |
| Angular Gyrus | Right | 35 | -61 | 44 | 4.420 |
| Cuneus | Left | -3 | -70 | 29 | 8.341 |
| Middle Cingulate | Left | -2 | -28 | 34 | 7.646 |
| Inferior Temporal | Left | -57 | -51 | -16 | 4.473 |
| Superior Frontal | Left | -17 | 17 | 64 | 4.694 |
| Postcentral Gyrus | Left | -47 | -42 | 58 | 4.453 |
| Unknown Control > Math |  |  |  |  |  |
| ParaHippocampal | Left | -29 | -31 | -19 | 6.541 |
| Middle Temporal | Left | -53 | 1 | -27 | 4.781 |

**Table S2.** MNI coordinates and t values of the main activation peaks for the contrasts of known > unknown math statements, unknown > known math statements, known > unknown control statements, and unknown > known control statements, extracted from maps thresholded at voxel-wise p < 0.001 uncorrected, and cluster-wise p < 0.05 FDR-corrected.

| Cortical Region | Hemisphere | MNI Peak Coordinates |  |  | t-value |
| --- | --- | --- | --- | --- | --- |
|  |  | x | y | z |  |
| Known > Unknown Math |  |  |  |  |  |
| Inferior Temporal | Left | -57 | -63 | -9 | 7.505 |
| Inferior Temporal | Right | 57 | -55 | -12 | 8.421 |
| Inferior Parietal | Left | -51 | -40 | 53 | 4.616 |
| Inferior Parietal | Right | 51 | -37 | 53 | 5.353 |
| Superior Parietal | Left | -15 | -76 | 52 | 5.108 |
| Superior Parietal | Right | 23 | -67 | 58 | 5.850 |
| Superior Frontal | Left | -21 | 7 | 70 | 5.730 |
| Superior Frontal | Right | 24 | -3 | 52 | 5.683 |
| Middle Frontal | Left | -32 | 29 | 49 | 5.007 |
| Middle Occipital | Left | -41 | -82 | 29 | 6.561 |
| Middle Occipital | Right | 45 | -78 | 28 | 7.092 |
| Anterior Cingulate | Left | -5 | 41 | -1 | 3.814 |
| Precuneus | Left | -9 | -61 | 64 | 5.212 |
| Unknown > Known Math |  |  |  |  |  |
| Precuneus | Left | -9 | -61 | 32 | 4.104 |
| Cuneus | Right | 12 | -70 | 35 | 7.028 |
| Superior Temporal | Left | -62 | -10 | -1 | 5.871 |
| Middle Temporal | Left | -68 | -27 | 7 | 5.943 |
| Superior Temporal | Right | 56 | -25 | 1 | 7.622 |
| Temporal Pole | Right | 57 | 7 | -7 | 6.509 |
| Inferior Frontal p. Tri | Left | -39 | 23 | 20 | 5.370 |
| Middle Occipital | Left | -32 | -82 | -1 | 4.959 |
| Middle Occipital | Right | 41 | -85 | 7 | 4.598 |
| Cerebelum 6 | Left | -29 | -60 | -21 | 4.317 |
| Cerebelum 6 | Right | 33 | -57 | -25 | 5.151 |
| Known > Unknown Control |  |  |  |  |  |
| Anterior Cingulate | Left | -8 | 55 | 2 | 5.478 |
| Unknown > Known Control |  |  |  |  |  |
| Superior Temporal | Left | -59 | -21 | 7 | 4.859 |
| Superior Temporal | Right | 57 | -19 | 5 | 3.333 |
| Middle Temporal | Left | -66 | -28 | 5 | 7.344 |
| Postcentral Gyrus | Left | -45 | -15 | 49 | 5.194 |
| Middle Cingulate | Right | 11 | 20 | 38 | 4.088 |
| Lingual Gyrus | Left | -20 | -55 | -3 | 4.859 |
| Lingual Gyrus | Right | 20 | -66 | -10 | 5.117 |

**Table S3.** MNI coordinates and t values of the main activation peaks for the contrasts of taught math statements in post-tests > pre-tests, taught control statements in post-tests > pre-tests, the interaction of taught > untaught statements in post-tests > pre-tests, and the interaction of taught > untaught and known statements in post-tests > pre-tests, extracted from maps thresholded at voxel-wise p < 0.001 uncorrected, and cluster-wise p < 0.05 FDR-corrected.

| Cortical Region | Hemisphere | MNI Peak Coordinates |  |  | t-value |
| --- | --- | --- | --- | --- | --- |
|  |  | x | y | z |  |
| Post > Pre Math Taught |  |  |  |  |  |
| Cuneus | Left | -11 | -67 | 28 | 7.683 |
| Cuneus | Right | 15 | -63 | 26 | 6.974 |
| Inferior Parietal Lobule | Left | -41 | -52 | 49 | 5.193 |
| Middle Frontal | Left | -39 | 52 | 8 | 5.245 |
| Inferior Frontal p. Tri | Left | -42 | 20 | 29 | 4.574 |
| Caudate Nucleus | Left | -11 | 17 | 5 | 4.702 |
| Caudate Nucleus | Right | 9 | 22 | -1 | 4.609 |
| Middle Occipital | Left | -38 | -78 | 40 | 4.756 |
| Post > Pre Control Taught |  |  |  |  |  |
| Cuneus | Left | -8 | -69 | 28 | 9.555 |
| Cuneus | Right | 15 | -64 | 28 | 9.280 |
| Angular Gyrus | Left | -41 | -73 | 38 | 5.483 |
| Inferior Parietal Lobule | Left | -48 | -45 | 49 | 3.480 |
| Angular Gyrus | Right | 44 | -70 | 37 | 4.304 |
| Middle Frontal | Left | -39 | 53 | 8 | 5.485 |
| Inferior Frontal p. Tri | Left | -42 | 22 | 31 | 5.354 |
| Caudate Nucleus | Left | -12 | 7 | 16 | 4.708 |
| Caudate Nucleus | Right | 18 | 10 | 17 | 4.208 |
| Middle Occipital | Left | -29 | -57 | 37 | 5.511 |
| Post > Pre x Taught > Untaught |  |  |  |  |  |
| Cuneus | Left | -14 | -64 | 23 | 8.568 |
| Cuneus | Right | 15 | -64 | 26 | 8.832 |
| Posterior Cingulate | Right | 6 | -45 | 11 | 4.789 |
| Inferior Parietal Lobule | Left | -41 | -54 | 49 | 4.959 |
| Angular Gyrus | Right | 41 | -70 | 37 | 4.407 |
| Middle Frontal | Left | -39 | 53 | 8 | 4.909 |
| Inferior Frontal p. Tri | Left | -42 | 22 | 31 | 5.614 |
| Caudate Nucleus | Left | -12 | 7 | 16 | 5.912 |
| Caudate Nucleus | Right | 9 | 20 | 1 | 5.276 |
| Middle Occipital | Left | -39 | -73 | 31 | 5.812 |
| Post > Pre x Taught > (Untaught + Known) |  |  |  |  |  |
| Cuneus | Left | -14 | -64 | 22 | 8.689 |
| Cuneus | Right | 15 | -64 | 26 | 8.523 |
| Inferior Parietal Lobule | Left | -41 | -54 | 49 | 5.039 |
| Angular Gyrus | Right | 42 | -72 | 35 | 4.586 |
| Superior Frontal | Left | -23 | 11 | 52 | 4.839 |
| Middle Frontal | Left | -39 | 52 | 7 | 4.807 |
| Inferior Frontal p. Tri | Left | -42 | 20 | 29 | 6.615 |
| Caudate Nucleus | Left | -12 | 7 | 16 | 7.090 |
| Caudate Nucleus | Right | 9 | 17 | 5 | 6.804 |
| Middle Occipital | Left | -38 | -75 | 31 | 6.926 |

**Table S4.** MNI coordinates and t values of the main activation peaks for the contrasts of greater intersubject correlations during math than during control video lessons, and vice-versa, extracted from maps thresholded at voxel-wise p < 0.001 uncorrected, and cluster-wise p < 0.05 FDR-corrected.

| Cortical Region | Hemisphere | MNI Peak Coordinates |  |  | t-value |
| --- | --- | --- | --- | --- | --- |
|  |  | x | y | z |  |
| ISC Math > Control Videos |  |  |  |  |  |
| Inferior Parietal Lobule | Left | -48 | -42 | 37 | 7.757 |
| Inferior Parietal Lobule | Right | 30 | -42 | 38 | 7.151 |
| Superior Partietal Lobule | Right | 51 | -40 | 58 | 7.133 |
| Inferior Temporal Gyrus | Left | -54 | -46 | -13 | 4.105 |
| Inferior Temporal Gyrus | Right | 51 | -42 | -15 | 5.656 |
| Superior Frontal Gyrus | Left | -23 | 1 | 55 | 7.064 |
| Superior Frontal Gyrus | Right | 26 | 7 | 61 | 9.142 |
| Middle Frontal Gyrus | Left | -39 | 25 | 37 | 6.368 |
| Middle Frontal Gyrus | Right | 44 | 35 | 32 | 7.218 |
| Inferior Frontal Gyrus | Left | -45 | 4 | 25 | 4.858 |
| Inferior Frontal Gyrus | Right | 45 | 7 | 23 | 6.945 |
| Cuneus | Left | -14 | -63 | 20 | 6.328 |
| Precuneus | Left | -17 | -66 | 50 | 6.507 |
| Precuneus | Right | 12 | -66 | 55 | 6.010 |
| ISC Control > Math Videos |  |  |  |  |  |
| Inferior Temporal Gyrus | Left | -42 | -31 | -21 | 3.235 |
| Lingual Gyrus | Left | -17 | -87 | -19 | 3.369 |
| Fusiform Gyrus | Left | -24 | -6 | -34 | 8.174 |
| Fusiform Gyrus | Right | 35 | -34 | -25 | 7.615 |
| Calcarine Sulcus | Right | 12 | -94 | -3 | 3.347 |
| Superior Temporal Gyrus | Left | -54 | -6 | -7 | 6.070 |
| Medial Temporal Pole | Left | -32 | 11 | -37 | 5.333 |
| Temporal Pole Superior | Right | 62 | -7 | -13 | 7.678 |
| Middle Temporal Gyrus | Left | -53 | -72 | 20 | 6.576 |
| Precentral Gyrus | Left | -51 | -12 | 37 | 5.029 |
| Precentral Gyrus | Right | 48 | -4 | 29 | 6.207 |
| Postcentral Gyrus | Right | 33 | -34 | 58 | 5.688 |
| Superior Frontal Gyrus | Left | -17 | 28 | 46 | 5.940 |
| Superior Frontal Gyrus | Right | 12 | 44 | 50 | 6.196 |
| Superior Frontal Gyrus | Right | 20 | 29 | 50 | 5.178 |
| IFG (p. Triangularis) | Left | -54 | 28 | 20 | 5.706 |
| Cerebelum (IX) | Left | -15 | -51 | -46 | 4.368 |
| Cerebelum (IX) | Right | 8 | -54 | -43 | 6.418 |
| Cerebelum (Crus 2) | Right | 12 | -85 | -40 | 5.103 |

